# A sulfatide-centered ultra-high resolution magnetic resonance MALDI imaging benchmark dataset for MS1-based lipid annotation tools

**DOI:** 10.1101/2025.10.20.683394

**Authors:** Lars Gruber, Stefan Schmidt, Thomas Enzlein, Carsten Hopf

## Abstract

Spatial ‘omics techniques are indispensable for studying complex biological systems and for the discovery of spatial biomarkers. While several current matrix-assisted laser desorption/ionization (MALDI) mass spectrometry imaging (MSI) instruments are capable of localizing numerous metabolites at high spatial and spectral resolution, the majority of MSI data is acquired at the MS1 level only. Assigning molecular identities based on MS1 data presents significant analytical and computational challenges, as the inherent limitations of MS1 data preclude confident annotations beyond the sum formula level. To enable future advancements of computational lipid annotation tools, well-characterized benchmark - or ground truth - datasets are crucial, which exceed the scope of synthetic data or data derived from mimetic tissue models. To this end, we provide two sulfatide-centered, biology-driven magnetic resonance MSI (MR-MSI) datasets at different mass resolving powers that characterize lipids in a mouse model of human metachromatic dystrophy. This data includes an ultra-high-resolution (R ∼1,230,000) quantum cascade laser mid-infrared imaging-guided MR-MSI dataset that enables isotopic fine structure analysis and therefore enhances the level of confidence substantially. To highlight the usefulness of the data, we compared 118 manual sulfatide annotations with the number of decoy database-controlled sulfatide annotations performed in Metaspace (67 at FDR < 10%). Overall, our datasets can be used to benchmark annotation algorithms, validate spatial biomarker discovery pipelines, and serve as a reference for future studies that explore sulfatide metabolism and its spatial regulation.

## DATA DESCRIPTION

The absence of ground truth datasets, i.e., prior knowledge of which metabolites/lipid are present (or not) in a tissue of interest, and the non-availability of corresponding datasets containing high-confidence annotations for a large number of metabolites/lipids, has been posing a major obstacle to computational advancements in mass spectrometry imaging (MSI) [1]. In particular, datasets that can challenge computational tools for molecular annotation will be crucial for rapid progress in the field [2]. To this end, we generated reusable and widely applicable datasets comprising quadruplicates of spatially focused, high-resolution mass spectrometry imaging data (MS1 level) derived from kidneys of an arylsulfatase A-deficient (ARSA-/-) mouse, a well-known genetic model of human metachromatic leukodystrophy [3]. Specifically, we developed a workflow that leverages Quantum Cascade Laser Mid-infrared (QCL-MIR) imaging to guide MSI on a 7T FT-ICR magnetic resonance mass spectrometer (MR-MS; **Supplementary Fig. 1 and 2**). The resulting MS1 data was interpreted in conjunction with precise reference annotations obtained for sulfatide glycosphingolipid species in defined kidney regions by on-tissue fragmentation-based lipid identification using imaging parallel reaction monitoring - parallel acquisition serial fragmentation (iprm-PASEF) on an orthogonal trapped ion mobility spectrometry (tims) TOF mass spectrometer [4]. Through combination of ultra-high resolution (R ∼ 1,230,000) MR-MSI MS1 data with systematic MS2 data obtained on a different mass spectrometer, we are establishing a concept for generating such benchmark datasets. Four biological replicates and cross-modal validation against 4D-lipidomics TIMS-MS ensure data quality [4]. The ultra-high resolution dataset was further intended to be complemented by a high-resolution dataset. All files and preprocessing scripts are publicly available, thus supporting benchmarking and integration within spatial omics analyses, as further demonstrated in this work using Metaspace-ML [5].

## CONTEXT

MALDI mass spectrometry imaging (MSI) has evolved into an invaluable tool in spatial biology [6, 7] that enables the label-free detection and statistically validated visualization of molecular distributions in tissues [8, 9]. However, achieving reliable bimolecular interpretation of the inherently complex spatial molecular patterns fundamentally depends on the availability of high-quality datasets featuring unambiguous molecular identifications. Such datasets are crucial for facilitating the discovery of spatial biomarkers and yielding insights into tissue function, pathology, and pharmacodynamic or therapeutic responses [10, 11].

Prompted by the instrumental limitations outlined, for instance, in the 4S paradigm [7], several specialized methodologies for subspace imaging have been developed to facilitate the generation of high-quality mass spectrometry imaging (MSI) datasets, including spatial sparse sampling strategies [12, 13] or guided approaches. The latter comprises mass-guided approaches, e.g., single-cell imaging [14] or on-tissue MS2 [15] and imaging-guided approaches [16–20], including the recently developed QCL-MIR imaging-guided MSI workflow [4]. In general, these workflows have been introduced with the objective of enhancing overall throughput. Sometimes, acquisition time saved by restricting MSI to defined ROIs is reallocated to alternative workflows that operate MSI with advanced instrumental settings for MS1 data. These alternative workflows can include adjustments to laser beam settings [21], increased transient durations in FT-ICR MSI, or optimized ramp times in TIMS-MSI, which improve spatial resolution, mass resolution, and ion mobility separation, respectively. Furthermore, imaging-based guidance methods can be combined with a sophisticated MSI technique for on-tissue MS2 that utilizes ion mobility-enhanced methods such as iprm-PASEF [4, 15]. This integration substantially increased data quality by enhancing the confidence level for molecular identifications, all without the need for high-performance liquid chromatography (HPLC) separation and directly in the spatial context of the tissue [22]. Notably, the exploration of the chemical space of sulfatide isoforms in an ARSA-/-mouse model enabled us to introduce a ground truth to the MSI field, since sulfatides are known to accumulate in distinct ROIs of kidney sections from these mice.

As our QCL-MIR imaging-based guidance approach is inherently instrument-agnostic, we have applied it in this study to create an ultra-high-resolution, sulfatide-focused MS1 benchmark dataset using a 7T XR FT-ICR (**Supplementary Fig. 1 and 2**). Using externally validated annotations, this dataset may become a unique benchmark resource (**Fig. 1a**) for developing and enhancing MS1 tools for deep spatial lipidomics and related research fields. This is especially important, because most MSI studies today still rely on acquiring only MS1 data [23, 24].

**Figure 1.**
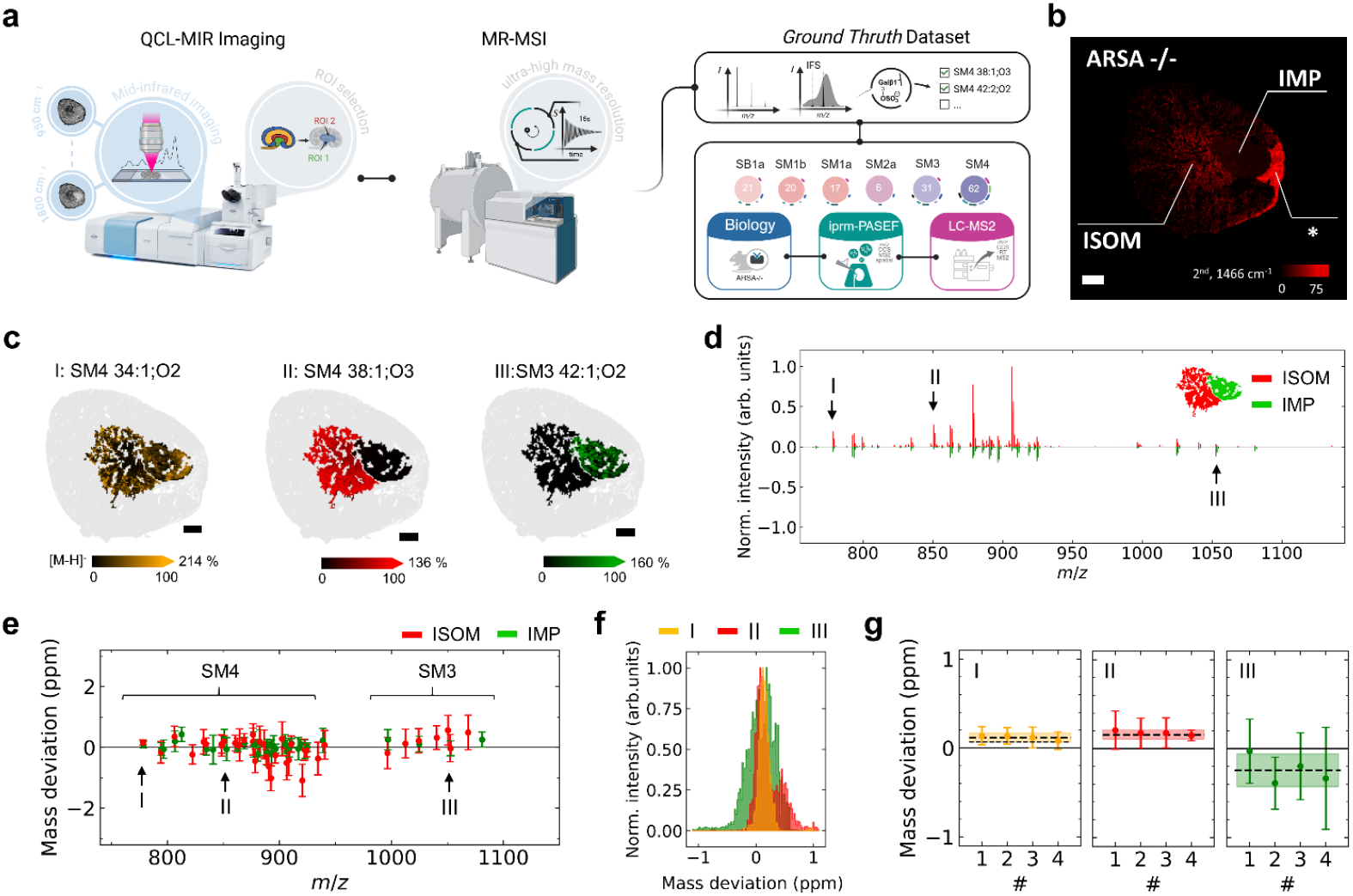
A sulfatide-centered, ultra-high resolution QCL-MIR guided MSI benchmark dataset. **a**, Schematic overview of data acquisition and incorporation of reference information. The methodology includes quantum cascade laser mid-infrared (QCL-MIR) imaging of kidney sections for region-focused magnetic resonance mass spectrometry imaging (MR-MSI) at ultra-long transient times (∼ 16s), followed by manual sulfatide annotations based on a reference dataset validated at MS2 level [4]. **b**, Representative lipid-distribution in an ARSA-/-kidney section based on the 2^nd^ derivative of absorbance (2^nd^) at 1466 cm^-1^ yielded predominant sulfatide accumulation in the ISOM region. The asterisk marks a region of high lipid content as described in [4]. Scale bar, 500 µm. **c**, Overlay of region-focused ion images of **(I)** *m/z* 778.5146 (SM4 34:1;O2[M-H]^-^; orange), **(II)** *m/z* 850.5721 (SM4 38:1;O3[M-H] ^-^; red), and **(III)** *m/z* 1052.6923 (SM3 42:1;O2[M-H] ^-^; green) in kidney (grey). Mass window, ±3 ppm. **d**, Representative butterfly plot of average mass spectra for the inner medulla/papilla (IMP; green) and ISOM (red) identified by QCL-MIR. **e**, Mass deviation and uncertainties (standard deviation) for 47 sulfatides (signals present in at least 50 pixels of either region IMP or ISOM) at R_2_∼1.23 M. *m/z* values are shifted by +0.2 (IMP) or −0.2 (ISOM) for visualization. **f**, Histogram of sum intensities in ISOM and IMP for (I), (II), and (III) measured with a mass resolution of ∼1.23M at *m/z* 800. **g**, Weighted mean mass deviation (dotted line) and uncertainty (*n*=4) presented as internal error^1^ (filled area) for (I), (II), and (III).

By default, annotations of MS1 data are limited to the sum formula level. For this and other reasons, the unambiguous annotation of sum formulae to MS1 data remains a non-trivial task, even with ultra-high mass resolving power (R > 500.000) spectra, due to the sheer chemical diversity within biological samples [2, 25, 26]. Consequently, a single precise mass measurement may correspond to multiple candidate sum formulae, thereby complicating definitive assignment, even when the isotopic fine structures (IFS) can be resolved. Nevertheless, the progressive exploitation of accurate mass, isotopic envelope, and IFS information [27–32] substantially increases the reliability of metabolite annotation in MR-MSI-based spatial ‘omics studies, a process in which computational tools play a crucial role by enabling automated annotation workflows. For future advancement of such tools in MSI, well-characterized benchmark datasets with high mass accuracy will be essential to enable robust validation and method development. This highlights the necessity and reuse potential of the dataset introduced in this study.

## METHODS

### Quantum-cascade laser mid-infrared (QCL-MIR) imaging of mouse kidneys

Animal studies involving ARSA-/-mice and cryo-sectioning of kidneys have been described before [4]. To ultimately focus ultra-high resolution MR-MSI data generation on defined kidney regions of interest (ROI) on an adjacent tissue section, we used QCL-MIR imaging for a pre-scan, followed by segmentation of hyperspectral QCL-MIR data to define ROIs [4]. These were then transferred to the MR-MSI instrument (**Supplementary Fig. 1 and 2**). To this end, QCL-MIR imaging data was recorded in sweep scan mode within a spectral range of 950–1800 cm^-1^ at a spectral sampling interval of 4 cm^-1^ on a Hyperion II ILIM (Bruker Optics, Ettlingen, Germany) equipped with a 3.5x objective. Subsequently, ROIs were generated and selected using *in-house* software (https://github.com/CeMOS-Mannheim/QCL_MIR_guided_MSI) based on spatial sulfatide distributions in ARSA-/-mouse kidneys, which predominantly occur in the Inner Medulla/Papillae (IMP) and Inner Stripe of Outer Medulla (ISOM). Specifically, these ROIs were then targeted for MR-MSI data acquisition with transient times of 15.7s. All measurements were repeated for n=4 biological replicates.

### Matrix spray-coating

10 mg/mL DHAP was dissolved in 70% ACN with 125 mM ammonium sulfate. After sonication, 0.1% TFA and 3 µM of SM4 35:1;O2 (100 µg/mL (= 157.41 µM) in MeOH/chloroform 2:1) as internal standard (IS) were added. Matrix was applied with an M5 TM-Sprayer (HTX Technologies, Chapel Hill, USA). Temperatures of the spray nozzle and tray were 75 °C and 35 °C, respectively. The spraying parameters were as follows: Spray Nozzle Velocity: 1200 mm/min; Flow Rate: 0.1 mL/min; No. of Passes: 10; Track Spacing: 2 mm; Pattern: HH; Pressure: 10 psi; Gas Low Rate: 2 L/min; Nozzle Height: 40 mm; Drying Time: 0s.

### Magnetic resonance mass spectrometry imaging (MR-MSI) data acquisition

Ultra-high resolution MSI data was acquired on a solariX 7T XR Fourier Transform Ion Cyclotron Resonance (FT-ICR) MS (Bruker Daltonics, Bremen, Germany), equipped with a smartbeam II 2 kHz laser and ftms control 2.3.0 software (Bruker Daltonics, Build 92). Mass spectra were acquired in negative ion mode (*m/z* range 401.29–2600) and data acquisition size of 8 M, resulting in a free induction decay (FID) time of 15.7s according to and a mass resolving power of 1,230,000 at *m/z* 800. Reducing the number of data points in the time domain to 512k resulted in an FID time of 0.98 s and a mass resolving power of R∼77,000. Ion optics settings were constant for all measurements: funnel RF amplitude (150 Vpp), source octopole (5 MHz, 350 Vpp), and collision cell voltage: 1.5 V, cell: 2 MHz, 1200 Vpp. The source DC optics were also constant for all measurements (capillary exit: −200 V, deflector plate: −220 V, funnel 1: −150 V, skimmer 1: −15 V), as well as the ParaCell parameters (transfer exit lens: 30 V, analyzer entrance: 10 V, sidekick: 0 V, side kick offset: 1.5 V, front/back trap plate: −3.4 V, back trap plate quench: 30 V). Sweet excitation power for ion detection was set to 14 %, and ion accumulation time was 0.05 s. The transfer optics were as follows: time of flight: 1 ms, frequency: 4 MHz, and RF amplitude: 350 Vpp. The laser parameters were laser power: 32 %, laser shots: 20, laser frequency: 200 Hz, and laser focus: medium, at a lateral step size of 40 µm.

### Mass spectrometry imaging data analysis

For state-of-the-art annotation with Metaspace [33], the v2 (Metaspace ML [5]) algorithm and an *m/z* tolerance of 2 ppm were utilized. The *imzML* files were exported from SCiLS Lab (Version 2024a Pro, Bruker Daltonics). For manual annotation, we compared the QCL-MIR-guided MR-MSI data against the ground truth data from [4]. For direct comparison of ultra-high- and high-resolution datasets, MS-based data segmentation was employed to enable subspace modeling. Bisecting k-means clustering of the MSI data from whole-tissue kidney sections was performed via SCiLS Lab to delineate anatomical regions, specifically the inner stripe of the outer medulla (ISOM) and the Inner Medulla/Papillae (IMP). These region outlines were subsequently used to generate a region-focused dataset, facilitating a more targeted comparative analysis.

## MATERIALS

All chemicals and solvents were of HPLC-MS grade. Conductive indium tin oxide (ITO)-coated glass slides were purchased from Diamond Coatings (West Midlands, UK). The MALDI matrix 2,5-dihydroxyacetophenone (DHAP) was purchased from Thermo Fisher Scientific (Waltham, Massachusetts, USA). Acetonitrile (ACN), ethanol (EtOH), LC-MS water, 2-propanol (IPA), and ammonium sulfate (AmS) were obtained from VWR Chemicals (Darmstadt, Germany). The sulfatide standard C_17_ mono-sulfo galactosyl(β) ceramide d18:1/17:0 (SM4 35:1;O2) was purchased from Avanti Polar Lipids (Birmingham, USA). Trifluoroacetic acid (TFA), Mayer’s hemalum solution, hydrochloric acid, sodium bicarbonate, magnesium sulfate, eosin Y-solution 0.5%, xylene, and eukitt were purchased from Merck KGaA.

### DATA VALIDATION AND QUALITY CONTROL

To enable the advancement of MS^1^-based molecular annotation tools for spatial biomarker discovery pipelines and other purposes, we acquired and in-depth-characterized two sulfatide-centered, biology-driven datasets derived from an arylsulfatase A mouse model at two different mass resolving powers. This includes an ultra-high mass resolution MR-MSI dataset (R ∼ 1,230,000) acquired using our recently introduced Quantum Cascade Laser (QCL) MIR guidance approach [4], in this study, combined with a 7T FT-ICR mass spectrometer. Briefly summarized, this workflow utilizes QCL-MIR imaging microscopy to rapidly acquire hyperspectral data from tissue samples, here, fresh-frozen kidney sections. Subsequent application of unsupervised segmentation algorithms, specifically k-means clustering, allows for the identification of biologically relevant regions of interest (ROIs) within the kidney tissue, particularly inner medulla/papilla (IMP) and inner stripe of outer medulla (ISOM), where sulfatides are enriched in ARSA −/-mice. Ultra-high-resolution MR-MSI was then performed in a ROI-targeted manner on these two tissue morphologies within kidneys from two 12-week-old and two 60-week-old ARSA-/-mice (**Fig. 1a; Supplementary Fig. 1** and **2**). The QCL-MIR imaging enabled focus on just two morphologies, enhancing both analytical depth and data acquisition efficiency, e.g., measurement time. As a reference and benchmark for the molecular content of these tissue areas and the total number of potential sulfatide identifications, we relied on our published reference data, which is based on three pillars: a known biological pathway leading to lipid-class specific accumulation of sulfatides, iprm-PASEF-derived molecular identifications in conjunction with 4D lipidomics LC-MS data [4] (**Supplementary Table 1**).

We compared conventional MR-MSI of whole kidney slices at a mass resolution of R_1_∼77,000 at _m/z_ 800 (1s free induction decay (FID) time; 14,331 pixels, 40 x 40 µm^2^ pixel size; 5 hours of data acquisition) in the FT-ICR with QCL-MIR-guided analysis (**Fig. 1b**) focused on the ISOM and IMP ROIs, which achieved R_2_∼1,230,000 at *m/z* 800 (16s FID time; 2,672 pixels, 40 x 40 µm^2^ pixel size; 11.6 hours of data acquisition), resulting in a 16-fold increase in mass resolving power (**Supplementary Fig. 3**). For the QCL-MIR-guided MR-MSI dataset, three sulfatide ion images are presented as examples that displayed similar intensities in both ROIs (**I**, *m/z* 778.5146 (SM4 34:1;O2[M-H]^-^)), higher intensity in ISOM (**II**, *m/z* 850.5721 (SM4 38:1;O3[M-H]^-^)), or higher intensity in IMP (**III**, m/z 1052.6923 [SM3 42:1;O2[M-H]-)). Unique molecular fingerprints were obtained per region (**Fig. 1d**). In both datasets, the mass deviation was constant across the *m/z* range (750-1100) and was consistently below 2 ppm, even when considering the uncertainties across n=4 biological replicates (**Fig. 2e; Supplementary Fig. 4**). The maximum mass deviation was about 1 ppm for the two less intense sulfatide ions **(I)** and (**III)** and less than 0.2 ppm for ion **(II)** at R_1_∼77,000, improving to below 0.2 ppm for **(I)** and (**II)** (around 0.5 ppm for (**III)**) with a mass resolution of R_2_∼1,230,000 (**Fig. 1f; Supplementary Fig. 4**). The reproducibility of our data is emphasized by comparing the mass deviation across n=4 biological replicates for ions (**I)-(III)**, all of which showed values below 1 ppm (**Fig. 1g**). Overall, the uncertainty of *m/z* values was reduced by a factor of 3-6 for data acquired at ultra-high resolution on an FT-ICR instrument.

**Figure 2.**
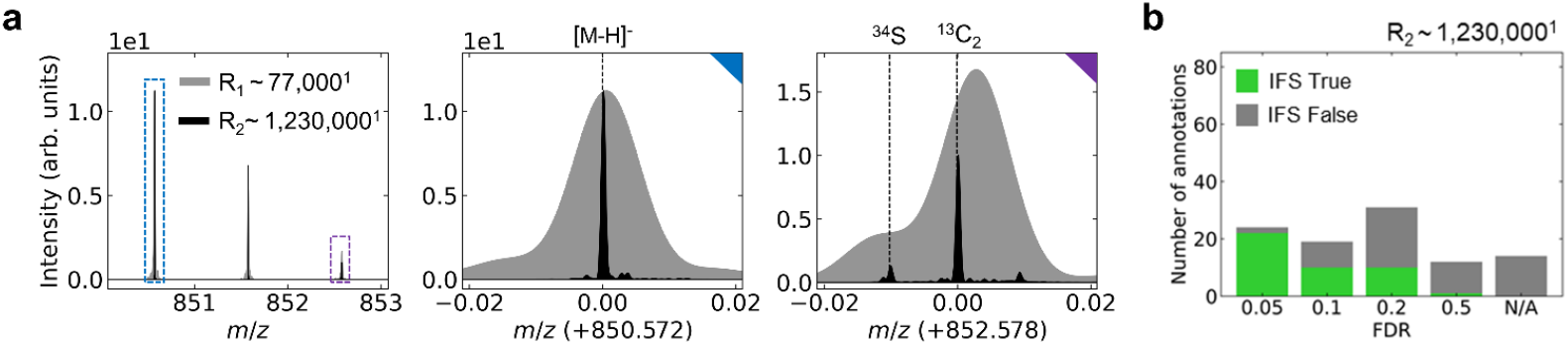
Evaluation of the annotation quality for the sulfatide-centered QCL-MIR guided MSI dataset. **a**, Ultra-high resolution MR-MSI data was acquired with a mass resolution of R_1_∼77,000 (gray) and R_2_∼1,230,000 (black). Isotopic fine structure (IFS) of SM4 38:1;O3[M-H]^-^ incl. ^13^C_2_ (M+2) and ^34^S isotopic peaks, normalized to the monoisotopic peak. **b**, Sulfatides were annotated using Metaspace, utilizing an in-house database consisting of LipidMaps fed with 780 theoretical sulfatides. ^1^Mass resolving power at *m/z* 800. N/A marks annotations that were performed manually (and validated via *in-situ* MS/MS and/or LC-MS/MS) but were not annotated by Metaspace at any FDR level

The ultra-high mass resolving power applied to the QCL-MIR dataset enables the detection of isotopic fine structures (IFS) for the sulfatides. The IFS, particularly the peak attributed to ^34^S, was very well resolved, with a signal-to-noise ratio of approximately 10 times the FWHM (**Fig. 2a; Supplementary Fig. 5; Supplementary Dataset 2**). Nevertheless, it is important to note that in all cases where FID times are notably prolonged, it is necessary to operate with a reduced total ion current. This precautionary measure is pivotal to reduce (local) space charge effects [34–37] within the ion cyclotron resonance (ICR) cell (**Supplementary Fig. 6**). However, this results in a loss of sensitivity, which in turn leads to slightly reduced numbers of sulfatide annotations, in particular for those isoforms that have a comparatively lower concentration in IMP and ISOM than in the cortex. Overall, the number of candidate sulfatide identifications using the QCL-MIR imaging-guided ultra-high-mass resolution MR-MSI method was 91 and 97 for the two 60-week ARSA-/-mice, compared to 118 and 115 annotations obtained with the conventional whole-tissue method (**Table 1**). The identification confidence was, on the other hand, dramatically improved, as 34 and 39 ultra-high-mass resolution spectra were supported by IFS information (**Table 1**). Identification was performed manually, as IFS is currently not utilized adequately in commercial sum formula annotation tools, which match experimental against theoretical isotope patterns [29]. In the sulfatide case, the ^34^S isotope peak was not automatically recognized as such, but pre-definition of S and N as constituents of the molecule-of-interest in the Bruker Smart Formula tool led to successful searches.

**Table 1:**
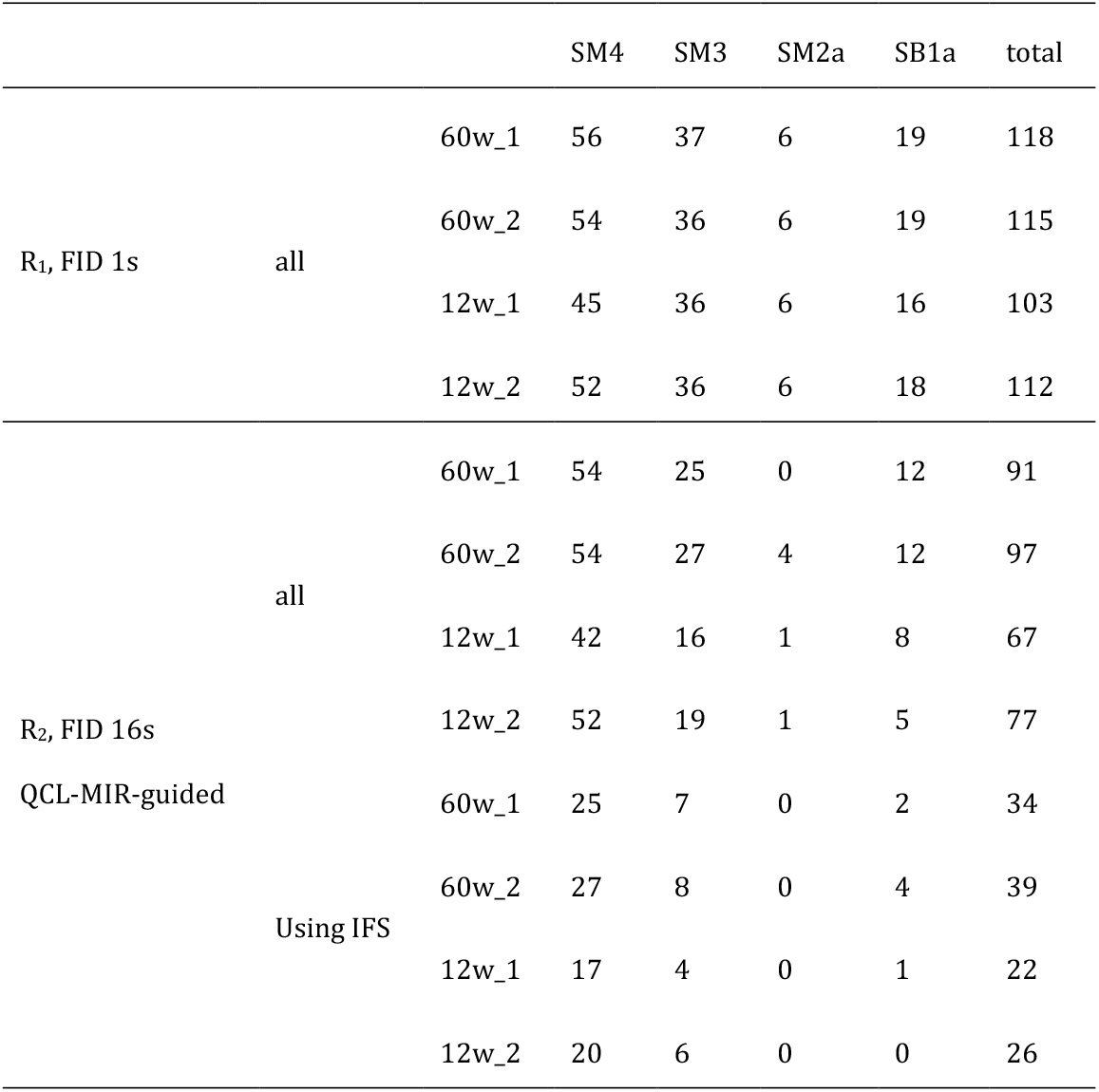
Cumulative numbers of sulfatide subclass isoforms identified in ARSA-/-mouse kidney by QCL-MIR imaging-guided MR-MSI. Whole kidney sections of 12- or 60-week-old ARSA-/-mice (n=2 each) were analyzed by conventional non-guided MR-MSI with a mass resolution of R_1_ ∼ 77k (at m/z 800) and QCL-MIR imaging-guided MR-MSI with a mass resolution of R_2_ ∼ 1,230k (at *m/z* 800). In many cases of QCL-MIR imaging-guided MSI, isotope fine structures (IFS) could be used for added confidence.

To highlight the potential reuse of our dataset as a benchmark for cutting-edge MALDI MSI data annotation, we used the open-source Metaspace platform (http://www.metaspace2020.eu). We compared the number of decoy database-controlled sulfatide annotations at different false-discovery rates (FDR). FDR-based quality measures have recently been critically assessed in the more mature field of proteomics [38], as i) FDR determination is often a black box, and ii) it is often unclear at what level FDR is set. For MSI, the FDR-control process in Metaspace is fairly transparent [33]. Since available databases do not yet sufficiently cover sulfatides, we manually created a database consisting of LipidMaps, which was additionally supplemented with 780 theoretical sulfatide structures (**Supplementary Dataset 3**). For data acquired at 77,000 mass resolving power, IMP and ISOM were extracted using MS feature-based segmentation to maintain consistency in the analysis. At the lowest FDR of 5%, we identified 24 sulfatide annotations for the 1.23M mass resolving power data, of which 22 were supported by IFS (**Fig. 2b**). In contrast, only 11 sulfatides were annotated in the 77k mass resolving power data (**Supplementary Fig. 7**). At 10% FDR, the ultra-high-resolution data yielded 43 annotations (32 supported by IFS), while the 77k mass resolving power data yielded 29 annotations. The number of unlikely annotations with FDR ≥20% was similar, with 74 (with 42 IFS; 1230k) and 77 (77k) sulfatide annotations, respectively. However, compared to our manual annotation, 14 (with 1 IFS; 1230k) and 19 (77k) sulfatides remained unannotated, even with an FDR of ≥50%. It should be noted that within the Metaspace annotation algorithm, only the four most intense peaks are recognized, regardless of whether an IFS is available. Overall, it is clear that even with high data quality, the current community standard tool, Metaspace, cannot eliminate the need for manual inspection of datasets.

### RE-USE POTENTIAL

This rigorously validated (see [4]) sulfatide-focused benchmark dataset offers substantial re-use potential for the MSI community. Acquired at two levels of mass resolving power, the dataset enables a comprehensive evaluation of annotation strategies, particularly those that leverage isotopic fine structure (IFS) for high-confidence MS1-level metabolite identification. As demonstrated, IFS analysis provides greater annotation accuracy than current automated platforms, underscoring the need for further methodological innovation in computational annotation. Although studies describing ultra-high resolution exist [39, 40], none of these studies shared their data under FAIR principles, and no externally validated annotations exist. This dataset serves as a valuable resource for benchmarking and validating spatial biomarker discovery pipelines, developing new annotation algorithms, and supporting future studies into sulfatide metabolism and its spatial regulation in biological tissues. Its broad applicability ensures relevance for both computational tool development and biological research, facilitating advances in spatial metabolomics and related fields.

## DATA AVAILABILITY

The underlying MALDI MR-MSI raw data supporting the findings of this study are openly available in Zenodo at http://doi.org/10.5281/zenodo.16842680. The.*imzML* files of the processed MR-MSI are available via Metaspace under https://metaspace2020.org/api_auth/review?prj=0e7b6e78-78cf-11f0-a049-172853cb2b10&token=EaH8S_2gCKoX.

## CODE AVAILABILITY

Data acquisition was conducted using existing tools (e,g, https://github.com/CeMOS-Mannheim/QCL_MIR_guided_MSI) and methods as described in the Methods section.

## List of abbreviations

ARSA: Arylsulfatase A
FDR: False discovery rate
FID: Free induction decay
FT-ICR: Fourier transform ion cyclotron resonance
FWHM: Full width at half maximum
IFS: Isotopic fine structure
IMP: Inner medulla/papillae
ISOM: Inner stripe of outer medulla
LC-MS: Liquid chromatography mass spectrometry
MALDI: Matrix-assisted laser desorption/ionization
MIR: Mid-infrared
MRMS: Magnetic resonance mass spectrometry
MSI: Mass spectrometry imaging
m/z: mass to charge ratio
QCL: Quantum cascade laser
ROI: Region of interest

## DECLARATIONS

### Author Contributions (CRediT)

Lars Gruber: Methodology, Investigation, Formal Analysis, Writing - Original Draft Stefan Schmidt: Methodology, Investigation, Formal Analysis, Writing - Original Draft Thomas Enzlein: Formal Analysis, Visualization Carsten Hopf: Conceptualization, Supervision, Project Administration, Resources, Writing - Review & Editing

### Funding (FundRef)

This work was supported by:

- The Bundesministerium für Bildung und Forschung (BMBF) under grants 12FH8I05IA (Drugs4Future) and 13FH8I09IA (DrugsData) within the M2Aind partnership (to Carsten Hopf);
- The Ministerium für Wissenschaft, Forschung und Kunst Baden-Württemberg (MWK) via the Mittelbauprogramm (to Carsten Hopf);
- The Deutsche Forschungsgemeinschaft (DFG, project 262133997) for the acquisition of the solariX 7T XR (to Carsten Hopf).

The funders had no role in study design, data collection/analysis, interpretation, or manuscript preparation.

## DECLARATION OF GENERATIVE AI AND AI −ASSISTED TECHNOLOGIES IN THE WRITING PROCESS

During the preparation of this work, the author(s) used perplexity.ai to improve the readability and language of the manuscript. After using this tool/service, the author(s) reviewed and edited the content as needed and take(s) full responsibility for the content of the published article.

## ASSOCIATED CONTENT

### Supporting Information

The Supporting Information is available free of charge.

