## Supplementary Information for "A sulfatide-centered ultra-high resolution magnetic resonance MALDI imaging benchmark dataset for MS1-based lipid annotation tools"

### SUPPLEMENTARY DATA

**Supplementary Dataset 1.** Evaluation of sulfatides peak analysis for conventional 7T XR FT-ICR data with 1s FID time.

**Supplementary Dataset 2.** Evaluation of sulfatides peak analysis for conventional 7T XR FT-ICR data with 16s FID time.

For data visualization and rapid data interpretation, ion images corresponding to a sulfatide feature list consisting of 126 features were exported from SCILS lab software. Additionally, centroided data derived from mean ion intensities and mass resolving power per peak were exported from Data Analysis software. Each peak was modeled using a Gaussian approximation. Data processing in Python identified the peak position of the monoisotopic mass, $^{13}C_2$, and $^{34}S$.

**Supplementary Dataset 3.** List of theoretical sulfatide configurations provided as .xlsx file.

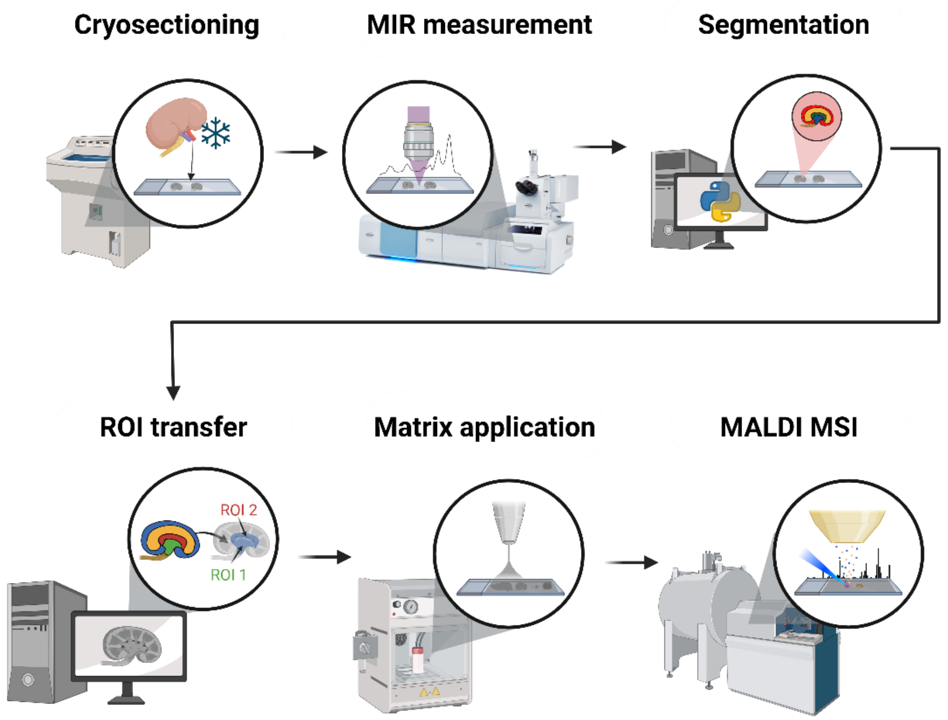


**Supplementary Figure 1. Schematic overview of the QCL-MIR imaging guidance and MR-MSI data acquisition workflow.**

Tissues (here, kidneys from arylsulfatase A knock-out mice) are cryosectioned and dried and scanned by quantum-cascade laser (QCL) mid-infrared (MIR) imaging microscopy. Hyperspectral datasets are segmented based on chemical composition of the tissue, and regions of interest (ROI) are transferred to an FT-ICR magnetic resonance mass spectrometry imaging (MR-MSI) instrument. The tissue section is spray-coated with a chemical MALDI matrix and subjected to MALDI MS imaging on the MR-MSI system at either high resolution (R ~ 77,000) or ultra-high resolution (R ~ 1,230,000).

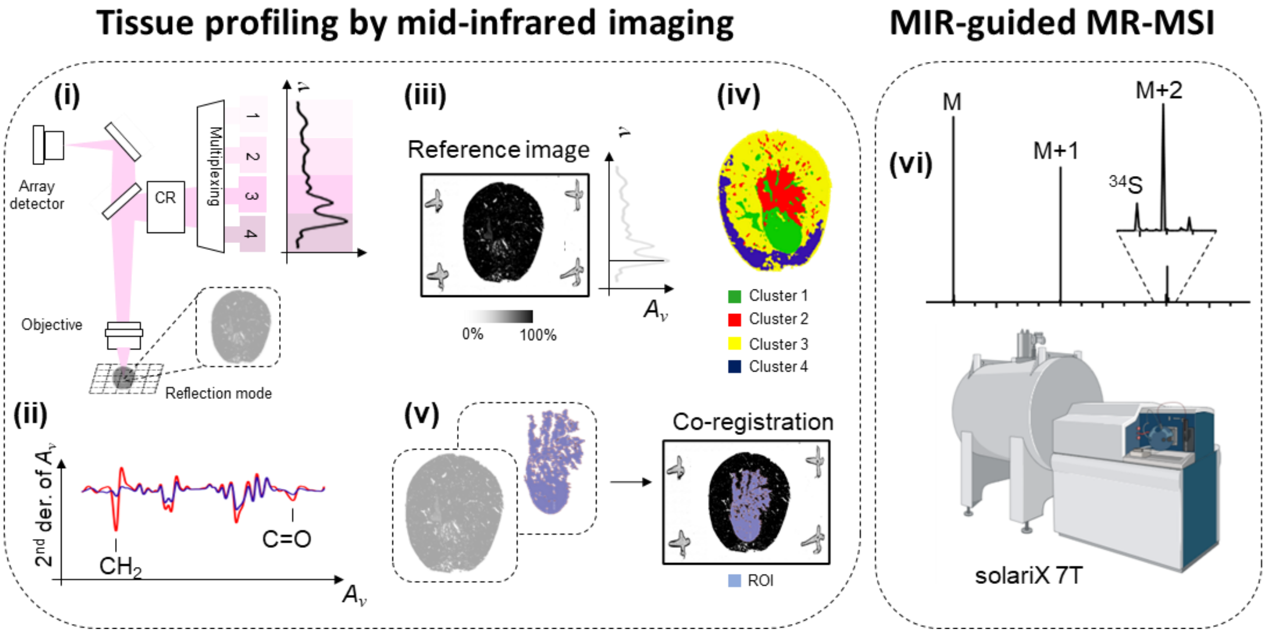


**Supplementary Figure 2. Detailed Schematic of the guidance and data acquisition workflow.**
Schematic overview of the QCL microscope (i) and the QCL-IRI-guided MSI workflow for fresh-frozen biological specimens in reflection mode. (ii) Key spectral features are selected using the second derivative of absorbance (1466 $cm^{-1}$, $CH_2$ bending; 1742 $cm^{-1}$, C=O stretch). (iii) A single wavenumber (1656 $cm^{-1}$) image is acquired as a reference and co-registered with the QCL-MIR dataset-derived regions-of-interest (iv and v). (vi) MR-MSI is performed with a focus on these ROIs, enabling the detection of isotopic fine structures of sulfatides. $A_\nu$: absorbance; CR: coherence reduction.

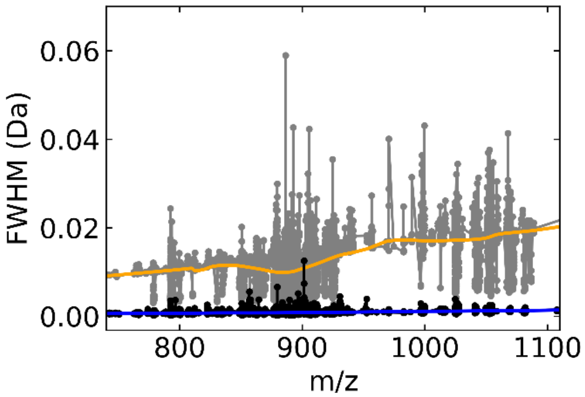


**Supplementary Figure 3. Evolution of experimentally deducted FWHM for two different mass resolving powers across the lipid mass range for datasets of a 60-week-old mouse kidney.**
Full width at half maximum (FWHM) as a function of the *m/z* value for two different free induction decay (FID) times of 1s (grey, mass resolving power $R_1$) and 16s (black, mass resolving power $R_2$). The orange (1s FID) and blue (16s FID) curves result from a locally estimated scatterplot smoothing (loess) and are plotted to guide the eye. On average, the ratio of the FWHM with $R_2$ to the FWHM with $R_1$ agrees well with the expected mass resolving power determined by the relative duration of the FID times.

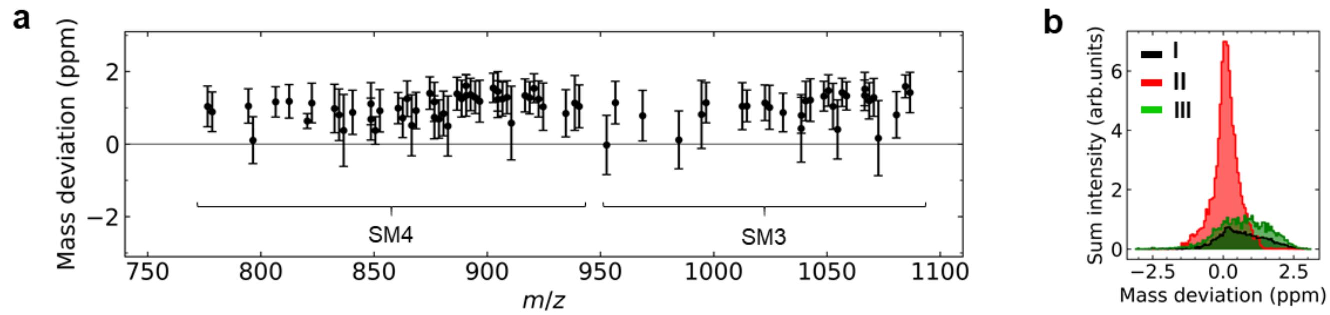


**Supplementary Figure 4. Mass deviation across the lipid mass range for $R_1$~77,000 across n=4 biological replicates of ARSA-/- mouse kidney.**
**a**, Mean mass deviation for sulfatides identified by MR-MSI (solariX 7T XR) with a mass resolution of $R_1$~77,000 at *m/z* 800. Means of mass deviation and standard deviations for each sulfatide were obtained from the sum intensity histograms in b, which resulted from the mean *m/z* value (determined across all pixels) of a given sulfatide. **b**, Histogram of the sum intensity for the three sulfatides I SM4 34:1;O2[M-H]$^-$, II SM4 38:1;O3[M-H]$^-$, and III SM3 42:1;O2[M-H]$^-$.

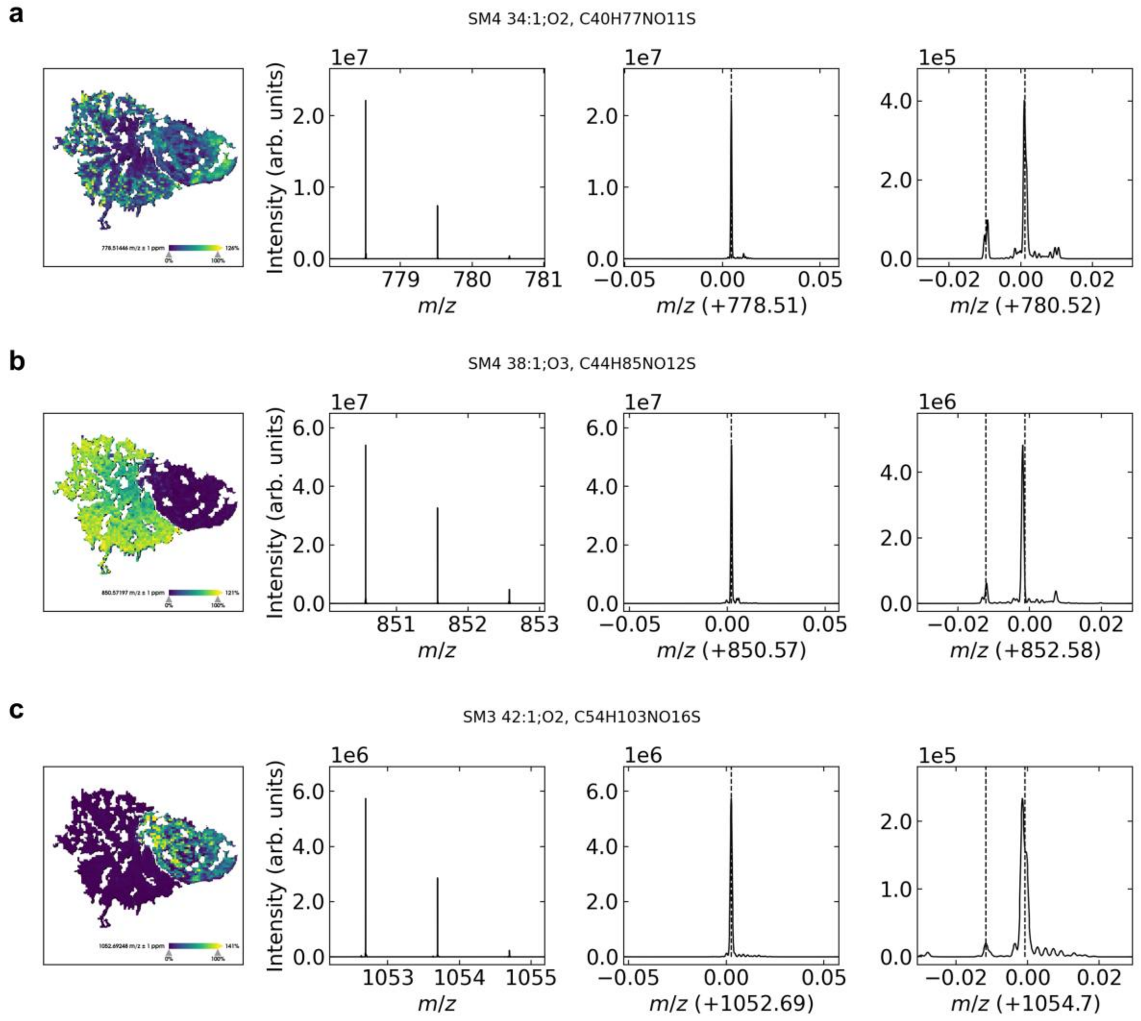


**Supplementary Figure 5. Spatial distribution of three sulfatide species and their respective isotopic fine structure.**
**a**, Ultra-high resolution MRMS data for I *m/z* 778.5146 (SM4 34:1;O2[M-H]$^-$; **b**, for II *m/z* 850.5721 (SM4 38:1;O3[M-H]$^-$; **c**, and III *m/z* 1052.6923 (SM3 42:1;O2[M-H]$^-$, acquired with a mass resolution of $R_2$~1,230,000 using QCL-MIR imaging-guided

MR-MSI. Positions of the isotopic fine structure (IFS) peak, including $^{13}C_2$ (M+2) and $^{34}S$, are highlighted by dotted lines, respectively. Respective data for all other sulfatides are presented in the Supplementary Dataset 2. Pixel size, 40 µm.

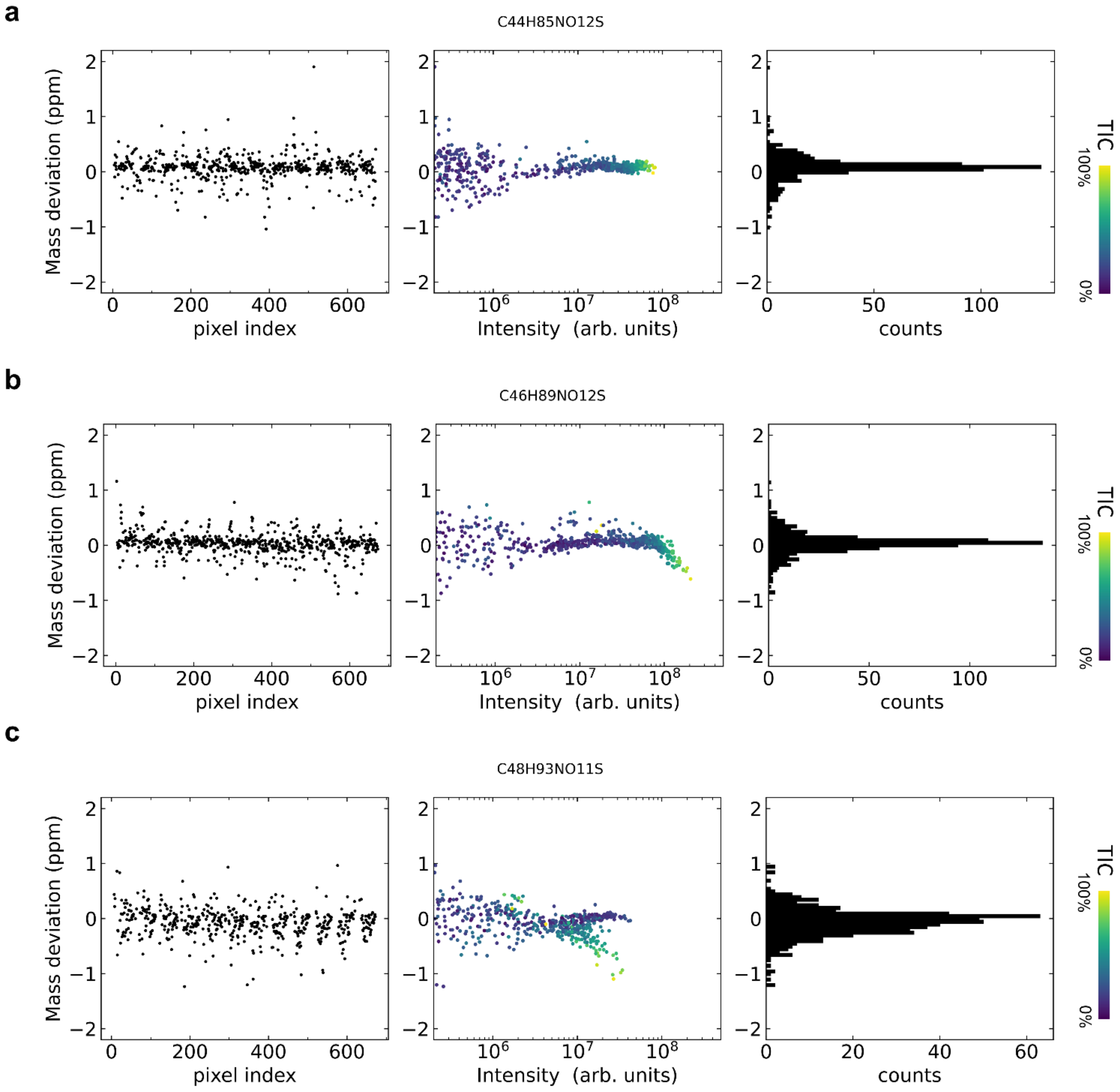


**Supplementary Figure 6. Observation of space-charge effects for three sulfatide examples.**
**a**, Evaluation for *m/z* 850.5720 (SM4 38:1;O3$[M-H]^-$; **b**, Evaluation for *m/z* 878.6033 (SM4 40:1;O3$[M-H]^-$; and **c**, Evaluation for *m/z* 890.6397 (SM4 42:1;O2$[M-H]^-$. All data was acquired with a mass resolution of $R_2$~1,230,000 using QCL-MIR imaging-guided MR-MSI. Bin width for histogram, 0.05 ppm.

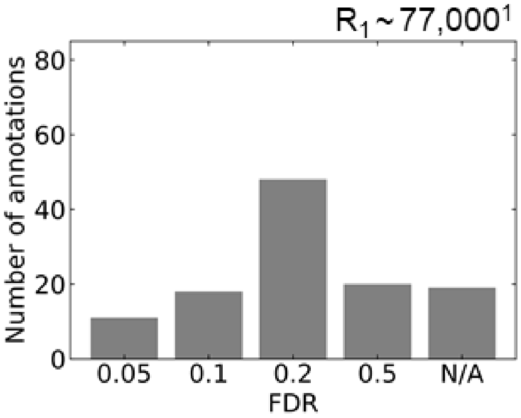


**Supplementary Figure 7. Evaluation of annotation quality for ARSA-/- kidney IMP/ISOM of the data with a mass resolving power of $R_1$.**
The annotation was performed via Metaspace utilizing an in-house database consisting of LipidMaps fed with 780 theoretical sulfatides. [1] at *m/z* 800. N/A marks annotations that were performed manually (and validated via in-situ MS/MS and/or LC-MS/MS) but were not annotated by Metaspace on any FDR level. [1]Mass resolving power at *m/z* 800.

**Supplementary Table 1. Overview of sulfatides identified in ARSA-/- kidney by MALDI-MSI.**
Uncertainties as standard deviation (*n*=4) in parentheses. Sulfatides with predominant accumulation in the cortex region are marked in italics. Sulfatides uniquely found in (Gruber *et al.* 2025) are marked with an asterisk and internal standard is marked with a hash mark. SM1a/b sulfatides were not considered here, since their [M-H]$^-$ can't be discriminated from SB1a [M-HSO3]$^-$, for details see (Gruber *et al.* 2025).

| Name | Sum formula | m/z theo. | Adduct | *m/z* FTICR | ppm FTICR |
|---|---|---|---|---|---|
| SM4 18:1;O2 | C24H47NO10S | 540.284791 | [M-H]$^-$ | 540.28479 | -0.097 |
| *SM4 18:0;O3 | C24H49NO11S | 558.295356 | [M-H]$^-$ | | |
| *SM4 32:2;O2 | C38H71NO11S | 748.467507 | [M-H]$^-$ | | |
| *SM4 32:1;O2 | C38H73NO11S | 750.483157 | [M-H]$^-$ | | |
| *SM4 32:2;O3 | C38H71NO12S | 764.462422 | [M-H]$^-$ | | |
| *SM4 32:1;O3 | C38H73NO12S | 766.478072 | [M-H]$^-$ | | |
| SM4 34:2;O2 | C40H75NO11S | 776.498807 | [M-H]$^-$ | 776.49881 | 0.061 |
| *SM4 33:2;O3 | C39H73NO12S | 778.478072 | [M-H]$^-$ | | |
| SM4 34:1;O2 | C40H77NO11S | 778.514457 | [M-H]$^-$ | 778.51446 | 0.116 |
| *SM4 34:2;O3 | C40H75NO12S | 792.493722 | [M-H]$^-$ | | |
| #SM4 35:1;O2 | C41H79NO11S | 792.530107 | [M-H]$^-$ | | |
| SM4 34:1;O3 | C40H77NO12S | 794.509372 | [M-H]$^-$ | 794.50937 | 0.006 |
| SM4 34:0;O3 | C40H79NO12S | 796.525022 | [M-H]$^-$ | 796.52502 | 0.008 |
| SM4 36:2;O2 | C42H79NO11S | 804.530107 | [M-H]$^-$ | 804.53011 | -0.079 |
| SM4 36:1;O2 | C42H81NO11S | 806.545757 | [M-H]$^-$ | 806.54576 | 0.124 |
| SM4 34:0;O4 | C40H79NO13S | 812.519936 | [M-H]$^-$ | 812.51994 | 0.123 |
| SM4 36:2;O3 | C42H79NO12S | 820.525022 | [M-H]$^-$ | 820.52502 | 0.077 |
| *SM4 37:1;O2 | C43H83NO11S | 820.561407 | [M-H]$^-$ | | |
| SM4 36:1;O3 | C42H81NO12S | 822.540672 | [M-H]$^-$ | 822.54067 | -0.056 |
| *SM4 36:0;O3 | C42H83NO12S | 824.556322 | [M-H]$^-$ | | |
| SM4 38:2;O2 | C44H83NO11S | 832.561407 | [M-H]$^-$ | 832.56141 | 0.084 |
| SM4 38:1;O2 | C44H85NO11S | 834.577057 | [M-H]$^-$ | 834.57706 | 0.051 |
| SM4 37:1;O3 | C43H83NO12S | 836.556322 | [M-H]$^-$ | 836.55632 | 0.006 |
| SM4 36:0;O4 | C42H83NO13S | 840.551237 | [M-H]$^-$ | 840.55124 | -0.04 |
| *SM4 39:2;O2 | C45H85NO11S | 846.577057 | [M-H]$^-$ | | |
| SM4 38:2;O3 | C44H83NO12S | 848.556322 | [M-H]$^-$ | 848.55632 | 0.075 |
| SM4 39:1;O2 | C45H87NO11S | 848.592707 | [M-H]$^-$ | 848.59271 | 0.066 |
| SM4 38:1;O3 | C44H85NO12S | 850.571972 | [M-H]$^-$ | 850.57197 | 0.154 |
| *SM4 38:0;O3 | C44H87NO12S | 852.587622 | [M-H]$^-$ | | |
| SM4 40:2;O2 | C46H87NO11S | 860.592707 | [M-H]$^-$ | 860.59271 | 0.118 |

| | | | | | |
|---|---|---|---|---|---|
| SM4 40:1;O2 | C46H89NO11S | 862.608357 | $[M-H]^-$ | 862.60836 | 0.057 |
| SM4 39:1;O3 | C45H87NO12S | 864.587622 | $[M-H]^-$ | 864.58762 | 0.041 |
| *SM4 39:0;O3 | C45H89NO12S | 866.603272 | $[M-H]^-$ | | |
| SM4 38:0;O4 | C44H87NO13S | 868.582536 | $[M-H]^-$ | 868.58254 | 0.132 |
| SM4 41:2;O2 | C47H89NO11S | 874.608357 | $[M-H]^-$ | 874.60836 | 0.134 |
| SM4 40:2;O3 | C46H87NO12S | 876.587622 | $[M-H]^-$ | 876.58762 | 0.162 |
| SM4 41:1;O2 | C47H91NO11S | 876.624007 | $[M-H]^-$ | 876.62401 | 0.063 |
| SM4 40:1;O3 | C46H89NO12S | 878.603272 | $[M-H]^-$ | 878.60327 | -0.077 |
| *SM4 40:0;O3 | C46H91NO12S | 880.618922 | $[M-H]^-$ | | |
| *SM4 39:0;O4 | C45H89NO13S | 882.598187 | $[M-H]^-$ | | |
| SM4 42:3;O2 | C48H89NO11S | 886.608357 | $[M-H]^-$ | 886.60836 | 0.013 |
| SM4 42:2;O2 | C48H91NO11S | 888.624007 | $[M-H]^-$ | 888.62401 | -0.122 |
| SM4 41:2;O3 | C47H89NO12S | 890.603272 | $[M-H]^-$ | 890.60327 | -0.035 |
| SM4 42:1;O2 | C48H93NO11S | 890.639657 | $[M-H]^-$ | 890.63966 | -0.103 |
| SM4 41:1;O3 | C47H91NO12S | 892.618922 | $[M-H]^-$ | 892.61892 | -0.235 |
| *SM4 41:0;O3 | C47H93NO12S | 894.634572 | $[M-H]^-$ | | |
| SM4 40:0;O4 | C46H91NO13S | 896.613837 | $[M-H]^-$ | 896.61384 | -0.119 |
| SM4 42:3;O3 | C48H89NO12S | 902.603272 | $[M-H]^-$ | 902.60327 | 0.083 |
| SM4 42:2;O3 | C48H91NO12S | 904.618922 | $[M-H]^-$ | 904.61892 | 0.149 |
| SM4 43:1;O2 | C49H95NO11S | 904.655307 | $[M-H]^-$ | 904.65531 | 0.032 |
| SM4 42:1;O3 | C48H93NO12S | 906.634572 | $[M-H]^-$ | 906.63457 | -0.073 |
| *SM4 42:0;O3 | C48H95NO12S | 908.650222 | $[M-H]^-$ | | |
| *SM4 41:0;O4 | C47H93NO13S | 910.629487 | $[M-H]^-$ | | |
| *SM4 44:3;O2 | C50H93NO11S | 914.639657 | $[M-H]^-$ | | |
| SM4 44:2;O2 | C50H95NO11S | 916.655307 | $[M-H]^-$ | 916.65531 | -0.033 |
| SM4 44:1;O2 | C50H97NO11S | 918.670957 | $[M-H]^-$ | 918.67096 | 0.087 |
| SM4 43:1;O3 | C49H95NO12S | 920.650222 | $[M-H]^-$ | 920.65022 | -0.034 |
| SM4 42:1;O4 | C48H93NO13S | 922.629487 | $[M-H]^-$ | 922.62949 | 0.007 |
| SM4 42:0;O4 | C48H95NO13S | 924.645137 | $[M-H]^-$ | 924.64514 | -0.046 |
| *SM4 44:2;O3 | C50H95NO12S | 932.650222 | $[M-H]^-$ | | |
| SM4 44:1;O3 | C50H97NO12S | 934.665872 | $[M-H]^-$ | 934.66587 | -0.085 |
| *SM4 43:0;O4 | C49H97NO13S | 938.660787 | $[M-H]^-$ | | |
| SM4 46:1;O2 | C52H101NO11S | 946.702258 | $[M-H]^-$ | 946.70226 | 0.323 |
| *SM4 44:0;O4 | C50H99NO13S | 952.676438 | $[M-H]^-$ | | |
| *SM3 18:1;O2 | C30H57N1O15S1 | 702.337615 | $[M-H]^-$ | | |
| *SM3 18:0;O3 | C30H59N1O16S1 | 720.348180 | $[M-H]^-$ | | |
| SM3 34:1;O2 | C46H87N1O16S1 | 940.567281 | $[M-H]^-$ | 940.56728 | 0.100 |
| SM3 34:1;O3 | C46H87N1O17S1 | 956.562196 | $[M-H]^-$ | 956.56220 | 0.113 |
| SM3 36:1;O2 | C48H91N1O16S1 | 968.598581 | $[M-H]^-$ | 968.59858 | -0.054 |
| *SM3 36:1;O3 | C48H91N1O17S1 | 984.593496 | $[M-H]^-$ | | |
| *SM3 38:2;O2 | C50H93N1O16S1 | 994.614231 | $[M-H]^-$ | | |
| SM3 38:1;O2 | C50H95N1O16S1 | 996.629881 | $[M-H]^-$ | 996.62988 | 0.053 |
| *SM3 38:2;O3 | C50H93N1O17S1 | 1010.609146 | $[M-H]^-$ | | |
| *SM3 39:1;O2 | C51H97N1O16S1 | 1010.645531 | $[M-H]^-$ | | |
| SM3 38:1;O3 | C50H95N1O17S1 | 1012.624796 | $[M-H]^-$ | 1012.62480 | 0.082 |
| SM3 38:0;O3 | C50H97N1O17S1 | 1014.640446 | $[M-H]^-$ | 1014.64044 | -0.226 |
| SM3 40:2;O2 | C52H97N1O16S1 | 1022.645531 | $[M-H]^-$ | 1022.64553 | 0.121 |
| SM3 40:1;O2 | C52H99N1O16S1 | 1024.661181 | $[M-H]^-$ | 1024.66118 | 0.068 |
| *SM3 38:0;O4 | C50H97N1O18S1 | 1030.635360 | $[M-H]^-$ | | |
| SM3 40:2;O3 | C52H97N1O17S1 | 1038.640446 | $[M-H]^-$ | 1038.64044 | -0.207 |
| SM3 41:1;O2 | C53H101N1O16S1 | 1038.676831 | $[M-H]^-$ | 1038.67683 | -0.099 |
| SM3 40:1;O3 | C52H99N1O17S1 | 1040.656096 | $[M-H]^-$ | 1040.65610 | 0.144 |
| SM3 40:0;O3 | C52H101N1O17S1 | 1042.671746 | $[M-H]^-$ | *1042.67175* | 0.036 |
| SM3 42:3;O2 | C54H99N1O16S1 | 1048.661181 | $[M-H]^-$ | 1048.66118 | -0.196 |
| SM3 42:2;O2 | C54H101N1O16S1 | 1050.676831 | $[M-H]^-$ | 1050.67683 | 0.163 |
| SM3 42:1;O2 | C54H103N1O16S1 | 1052.692481 | $[M-H]^-$ | 1052.69248 | -0.075 |
| SM3 41:1;O3 | C53H101N1O17S1 | 1054.671746 | $[M-H]^-$ | 1054.67175 | 0.101 |
| *SM3 41:0;O3 | C53H103N1O17S1 | 1056.687396 | $[M-H]^-$ | | |
| SM3 40:0;O4 | C52H101N1O18S1 | 1058.666661 | $[M-H]^-$ | 1058.66666 | 0.029 |
| *SM3 42:3;O3 | C54H99N1O17S1 | 1064.656096 | $[M-H]^-$ | | |
| SM3 42:2;O3 | C54H101N1O17S1 | 1066.671746 | $[M-H]^-$ | 1066.67175 | 0.103 |
| SM3 43:1;O2 | C55H105N1O16S1 | 1066.708131 | $[M-H]^-$ | 1066.70813 | 0.071 |
| SM3 42:1;O3 | C54H103N1O17S1 | 1068.687396 | $[M-H]^-$ | 1068.68740 | 0.146 |
| SM3 42:0;O3 | C54H105N1O17S1 | 1070.703046 | $[M-H]^-$ | *1070.70305* | 0.073 |
| *SM3 41:0;O4 | C53H103N1O18S1 | 1072.682311 | $[M-H]^-$ | | |
| *SM3 44:2;O2 | C56H105N1O16S1 | 1078.708131 | $[M-H]^-$ | | |
| SM3 44:1;O2 | C56H107N1O16S1 | 1080.723781 | $[M-H]^-$ | 1080.72378 | 0.111 |
| SM3 42:1;O4 | C54H103N1O18S1 | 1084.682311 | $[M-H]^-$ | 1084.68231 | 0.743 |
| SM3 42:0;O4 | C54H105N1O18S1 | 1086.697961 | $[M-H]^-$ | 1086.69796 | 0.101 |
| SM3 44:1;O3 | C56H107N1O17S1 | 1096.718696 | $[M-H]^-$ | 1096.71870 | 0.009 |

| | | | | | |
|---|---|---|---|---|---|
| *SM3 46:1;O2 | C58H111N1O16S1 | 1108.755082 | $[M-H]^-$ | | |
| *SM2a 38:1;O2 | C58H108N2O21S1 | 1199.709253 | $[M-H]^-$ | | |
| *SM2a 38:1;O3 | C58H108N2O22S1 | 1215.704168 | $[M-H]^-$ | | |
| *SM2a 40:1;O2 | C60H112N2O21S1 | 1227.740553 | $[M-H]^-$ | | |
| *SM2a 40:1;O3 | C60H112N2O22S1 | 1243.735468 | $[M-H]^-$ | | |
| *SM2a 42:1;O2 | C62H116N2O21S1 | 1255.771853 | $[M-H]^-$ | | |
| *SM2a 42:1;O3 | C62H116N2O22S1 | 1271.766768 | $[M-H]^-$ | | |
| SB1a 34:1;O2 | C60H110N2O29S2 | 1305.699477 | $[M-HSO_3]^-$ | 1305.69948 | 0.167 |
| SB1a 34:1;O3 | C60H110N2O30S2 | 1321.694392 | $[M-HSO_3]^-$ | 1321.69439 | -0.131 |
| SB1a 36:1;O2 | C62H114N2O29S2 | 1333.730777 | $[M-HSO_3]^-$ | 1333.73078 | 0.379 |
| SB1a 36:1;O3 | C62H114N2O30S2 | 1349.725692 | $[M-HSO_3]^-$ | 1349.72569 | 0.341 |
| SB1a 38:1;O2 | C64H118N2O29S2 | 1361.762077 | $[M-HSO_3]^-$ | 1361.76208 | 0.019 |
| SB1a 39:1;O2 | C65H120N2O29S2 | 1375.777727 | $[M-HSO_3]^-$ | 1375.77773 | 0.419 |
| *SB1a 38:1;O3 | C64H118N2O29S2 | 1377.756991 | $[M-HSO_3]^-$ | | |
| SB1a 40:2;O2 | C66H120N2O29S2 | 1387.777727 | $[M-HSO_3]^-$ | 1387.77773 | 0.505 |
| SB1a 40:1;O2 | C66H122N2O29S2 | 1389.793377 | $[M-HSO_3]^-$ | 1389.79338 | 0.170 |
| *SB1a 38:0;O4 | C64H120N2O31S2 | 1395.767556 | $[M-HSO_3]^-$ | | |
| SB1a 41:1;O2 | C67H124N2O29S2 | 1403.809027 | $[M-HSO_3]^-$ | 1403.80903 | 0.103 |
| SB1a 40:1;O3 | C66H122N2O30S2 | 1405.788292 | $[M-HSO_3]^-$ | 1405.78829 | -0.023 |
| SB1a 42:2;O2 | C68H124N2O29S2 | 1415.809027 | $[M-HSO_3]^-$ | 1415.80903 | -0.053 |
| SB1a 42:1;O2 | C68H126N2O29S2 | 1417.824677 | $[M-HSO_3]^-$ | 1417.82468 | 0.077 |
| *SB1a 40:0;O4 | C66H124N2O31S2 | 1423.798857 | $[M-HSO_3]^-$ | | |
| SB1a 42:2;O3 | C68H124N2O30S2 | 1431.803942 | $[M-HSO_3]^-$ | 1431.80394 | -0.120 |
| SB1a 42:1;O3 | C68H126N2O30S2 | 1433.819592 | $[M-HSO_3]^-$ | 1433.81959 | 0.087 |
| *SB1a 41:0;O4 | C67H126N2O31S2 | 1437.814507 | $[M-HSO_3]^-$ | | |
| *SB1a 44:1;O2 | C70H130N2O29S2 | 1445.855977 | $[M-HSO_3]^-$ | | |
| *SB1a 42:0;O4 | C68H128N2O31S2 | 1451.830157 | $[M-HSO_3]^-$ | | |
